# Androgens modulate the immune profile in a mouse model of polycystic ovary syndrome

**DOI:** 10.1101/2024.02.22.581579

**Authors:** Sara Torstensson, Angelo Ascani, Sanjiv Risal, Haojiang Lu, Allan Zhao, Alexander Espinosa, Eva Lindgren, Maria H. Johansson, Gustaw Eriksson, Maya Barakat, Mikael C.I. Karlsson, Camilla Svensson, Anna Benrick, Elisabet Stener-Victorin

## Abstract

Polycystic ovary syndrome (PCOS) is associated with a low-grade inflammation, but it is unknown how hyperandrogenism, the hallmark of PCOS, affects the immune system. Using a well-established PCOS-like mouse model, we demonstrate that androgen exposure affects immune cell populations in reproductive, metabolic, and immunological tissues differently in a site-specific manner. Co-treatment with flutamide, an androgen receptor antagonist, prevents most of these alterations, demonstrating that these effects are mediated through androgen receptor activation. Dihydrotestosterone (DHT)-exposed mice display a drastically reduced eosinophil population in uterus compared to controls, coupled with lower levels of eotaxin (CCL11), suggesting a reduced recruitment from blood. Decreased frequencies of eosinophils were also seen in visceral adipose tissue (VAT). A higher expression level of CD69, a marker of activation or tissue residency, was consistently found on natural killer (NK) cells in all analyzed tissues. However, a higher frequency of NK cells and elevated levels of IFN-γ and TNF-α were only seen in uteri of androgen-exposed mice, while NK cell frequencies were unaffected in all other analyzed compartments. Distinct alterations of macrophages in ovaries, uterus and VAT were also found in DHT-exposed mice and could potentially be linked to PCOS-like traits of the model. Indeed, androgen-exposed mice were insulin resistant and displayed an aberrant immune profile in VAT, albeit unaltered fat mass. Collectively, we demonstrate that hyperandrogenism causes tissue-specific alterations of immune cells in reproductive organs and VAT, which could have considerable implications on tissue function and contribute to the reduced fertility and metabolic comorbidities associated with PCOS.

## Introduction

The immune system is linked to several features of polycystic ovary syndrome (PCOS), including reproductive complications^1,2^ and metabolic comorbidities^3^. Yet, it remains largely unknown how hyperandrogenism, a hallmark of PCOS, affects the immune system.

Immune cells have essential functions in female reproductive organs, where uterine natural killer (NK) cells are crucial for successful implantation and the formation of endometrial spiral arteries^4^. Macrophages and other myeloid cells also play an important role during ovulation and mediate other essential ovarian functions, such as influencing sex hormone signaling in granulosa cells through the secretion of cytokines^5^. Endometrial dysfunction is associated with PCOS, indicated by altered expression of sex hormone receptors and abnormal regulation of enzymes involved in sex hormone biogenesis, dysregulated metabolic pathways and aberrant cell signaling^6^. An endometrial dysfunction is supported by a high prevalence of early miscarriage among women with PCOS, as well as pregnancy complications such as gestational diabetes, gestational hypertension and pre-eclampsia^1^. Similarly, immune cells in visceral adipose tissue are known to play a pivotal role in regulating metabolic function and adipose tissue inflammation is widely believed to be the underlying cause of insulin resistance and type 2 diabetes in obesity^7^. Substantial evidence shows that macrophages with a proinflammatory phenotype can cause insulin resistance^8^. Adipose tissue macrophages are regulated in a complex network where adipokines and proinflammatory cytokines, such as IFN-γ, trigger a more proinflammatory phenotype. Likewise, macrophages with an anti-inflammatory phenotype are essential to maintain insulin sensitivity in mice with diet-induced obesity^9^ and eosinophils have been shown to sustain these through the release of IL-4 and IL-13^10^. Hyperandrogenic women with PCOS have a higher risk of type 2 diabetes compared to women without PCOS, independent of BMI^3,11^. Furthermore, the prevalence of PCOS is higher among women with overweight or obesity compared to normal weight women^12^. Taken together, this could indicate that hyperandrogenism alters immune function, which causes a higher susceptibility to reproductive complications and metabolic comorbidities. Indeed, PCOS is associated with a low-grade inflammation^13–19^ but what impact this has on disease pathology and related comorbidities is unclear.

Due to practical limitations, human studies often fail to address the phenotypic and functional differences of circulating and tissue-resident immune cells. We therefore used a well-established PCOS-like mouse model^20^ to assess the impact of hyperandrogenism on the immune system in PCOS pathology. We determined the effect of dihydrotestosterone (DHT) exposure on major immune populations in reproductive, metabolic, and immunological tissues, and used co-treatment with flutamide, an androgen receptor (AR) blocker, to specifically study AR activation.

## Results

### Immune populations in blood and secondary lymphoid organs are affected by androgen exposure

To understand the systemic effect of hyperandrogenism on the immune system, we used flow cytometry to broadly analyze immune cell populations in blood and secondary lymphoid organs from peripubertal DHT-exposed PCOS-like mice (Fig. 1a and Supplementary Fig. 1a-b). An increased frequency of neutrophils (CD45^+^CD11b^+^SSC^mid^Ly6G^+^) and a reduced frequency of eosinophils (CD45^+^CD11b^+^SSC^high^Ly6G^-^Siglec-F^+^) was seen in blood of DHT-exposed mice compared to controls (Fig. 1b-c), an effect that was prevented by co-treatment with flutamide. The frequency of monocytes (CD45^+^CD11b^+^SSC^low^Ly6G^-^Ly6C^+^) was unchanged (Fig. 1d). To determine if the increased frequency of neutrophils was due to neutrophilia in DHT-exposed mice, the neutrophil counts were assessed. Surprisingly, there was no difference in neutrophil counts between the groups (Fig. 1e) while an overall decrease of CD45^+^ immune cells was observed in blood of DHT-exposed mice compared to controls and mice co-treated with flutamide (Fig. 1f). Concurrently, there was a trend suggesting a decreased frequency of T cells (CD45^+^CD3^+^) in DHT-exposed mice compared to those co-treated with flutamide (Fig. 1g). Subsequent analysis showed that T helper cells (CD45^+^CD3^+^CD4^+^) followed the same trend (Fig. 1h), while frequencies of cytotoxic T cells (CD45^+^CD3^+^CD8^+^) were unchanged among groups (Fig. 1i). These results could indicate an effect of androgen exposure on thymopoiesis, which was supported by a decreased thymic weight of DHT-exposed mice compared to controls and mice co-treated with flutamide (Fig. 1j). To determine the effect of androgens in secondary lymphoid organs, T cell populations were analyzed in lymph nodes and spleen. In contrast to the circulation, no effect was observed on T cell populations in spleen or lymph nodes following DHT-exposure (Supplementary Fig. 1c-j). The number of CD45^+^ immune cells was unchanged in spleen of DHT-exposed mice (Supplementary Fig. 1k), frequencies of immune cells should therefore correspond to absolute numbers. Moreover, the frequency of NK cells (CD45^+^CD3^-^NK1.1^+^) in blood was unaffected by androgen exposure (Fig. 1k). However, a higher proportion of NK cells expressed the activation marker CD69 in blood from DHT-exposed mice compared to controls (Fig. 1l) an effect that was not prevented by co-treatment with flutamide. Interestingly, the frequency of NK cells was higher in spleen of DHT-exposed mice compared to control and mice co-treated with flutamide (Fig. 1m). As in blood, the proportion of CD69^+^ NK cells was higher in spleen of DHT-exposed mice compared to controls (Fig. 1n) and was not prevented by co-treatment with flutamide. Taken together, androgen exposure affects immune populations differently in circulation and secondary lymphoid organs, with no uniform systemic effect.

**Figure 1.**
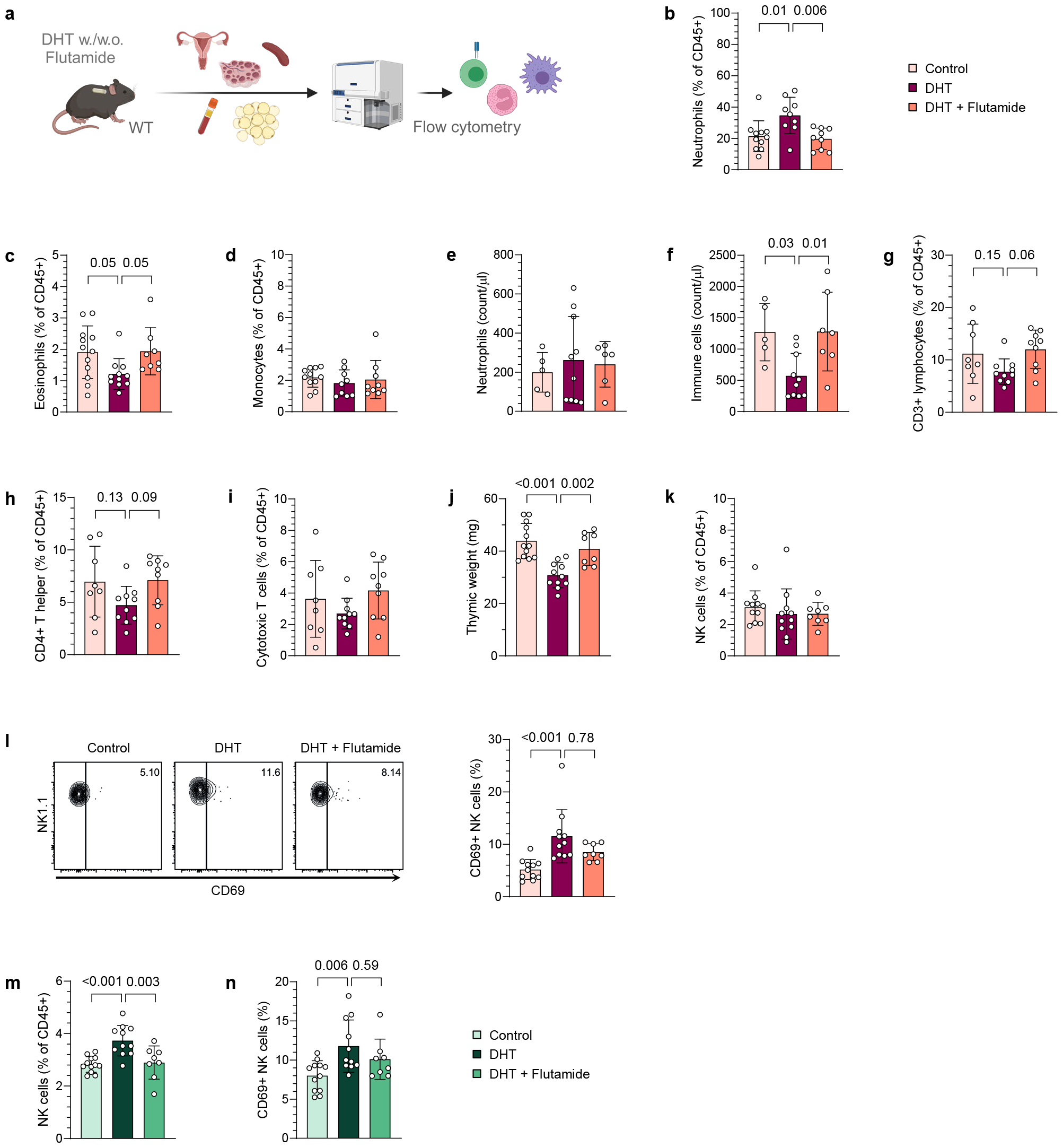
Immune populations in blood and secondary lymphoid organs are differently affected by androgen exposure. (**a**) Experimental design. (**b**) Frequency of neutrophils in blood, expressed as percent of CD45^+^ immune cells *(n* = 11 Control, 9 DHT, 9 DHT + Flutamide). (**c**) Frequency of eosinophils in blood *(n* = 12, 11, 8). (**d**) Frequency of monocytes in blood *(n* = 11, 9, 9). (**e**) Neutrophil count in blood *(n* = 5, 10, 6). (**f**) CD45+ immune cell count in blood *(n* = 5, 10, 7). (**g**) Frequency of CD3^+^ T cells in blood *(n* = 8, 10, 9). (**h**) Frequency of CD4^+^ T helper cells in blood *(n* = 8, 10, 9). (**i**) Frequency of CD8^+^ cytotoxic T cells in blood *(n* = 8, 10, 9). (**j**) Thymus weight *(n* = 12, 11, 8). (**k**) Frequency of NK cells in blood *(n* = 11, 11, 8). (**l**) CD69 expression on NK cells in blood *(n* = 11, 11, 8). (**m**) Frequency of NK cells in spleen *(n* = 12, 11, 8). (**n**) CD69 expression on NK cells in spleen *(n* = 12, 11, 8). Data are presented as means ± SD. n indicates the number of biologically independent samples examined. Statistical analysis was assessed by one-way ANOVA with Dunnett’s multiple comparison (**b, e**-**j, m**) or by Kruskal-Wallis with Dunn’s multiple comparison (**c**-**d, k**-**l, n**) and significant differences were indicated with p values. Source data are provided as a Source Data File.

### Uterine eosinophil and NK cell populations are markedly altered by androgen exposure in PCOS-like mice

Immune cells have essential functions in female reproductive organs, and endometrial dysfunction is associated with PCOS^6^. We therefore hypothesized that androgen exposure could contribute to endometrial dysfunction by altering the immune profile in the uterus. To test this hypothesis, major immune populations in uteri of DHT-exposed mice were first analyzed by flow cytometry (gating strategy in Supplementary Fig. 2a-b). Strikingly, the eosinophil population was drastically reduced in uteri of DHT-exposed mice compared to controls and mice co-treated with flutamide, which was seen as an evident decrease of the granular (SSC^high^) population (Fig. 2a-b). The NK cell population, on the other hand, was vastly increased in uteri of DHT-exposed mice, with no effect seen in mice co-treated with flutamide (Fig. 2c). DHT-exposure did not affect the absolute count of immune cells, and the decreased eosinophil population did not account for the observed increase of NK cells (Supplementary Fig. 2c-d). As in blood and other tissues, the proportion of NK cells expressing CD69 was higher in DHT-exposed mice compared to controls (Fig. 2d). Moreover, there was an increased frequency of CD45^+^CD11b^+^SSC^low^Ly6G^-^F4/80^+^ macrophages in uterus of DHT-exposed mice compared to controls and mice co-treated with flutamide (Fig. 2e). The increased frequency of macrophages did not appear to be due to a higher infiltration of monocytes as these were unaltered in uterus (Supplementary Fig. 2e). The macrophages in uteri of DHT-exposed mice also displayed a higher expression of MHC-II (Fig. 2f), suggesting a more pro-inflammatory phenotype. However, the expression of CD11c, considered a pro-inflammatory marker on macrophages, was unchanged (Fig. 2g). Next, uterine cytokine levels were assessed by Bio-Plex to define whether the effect on immune populations was associated with an altered cytokine profile. Eotaxin (CCL11) levels were lower in uterus of DHT-exposed mice (Fig. 2h), while levels of IL-5 were unchanged (Fig. 2i), suggesting that the reduced eosinophil population could be due to a disrupted recruitment from the periphery. In line with a higher activation state of NK cells, and a more pro-inflammatory phenotype of macrophages, levels of IFN-γ and TNF-α were higher in DHT-exposed mice, an effect that was prevented by co-treatment of flutamide (Fig. 2j-k). In agreement with the unaltered frequency on monocytes, the levels of CCL2 were unchanged among groups (Fig. 2l). Furthermore, uterine levels of IL-6 and granulocyte-colony stimulating factor (G-CSF) differed between DHT-exposed mice and mice co-treated with flutamide, but no significant difference was seen compared to controls (Supplementary Fig. 2f-g). No difference was seen among other analyzed cytokines (Supplementary Table 1). In summary, androgens has a drastic effect on uterine eosinophils and NK cells and related cytokines, an effect that is mediated through AR activation.

**Figure 2.**
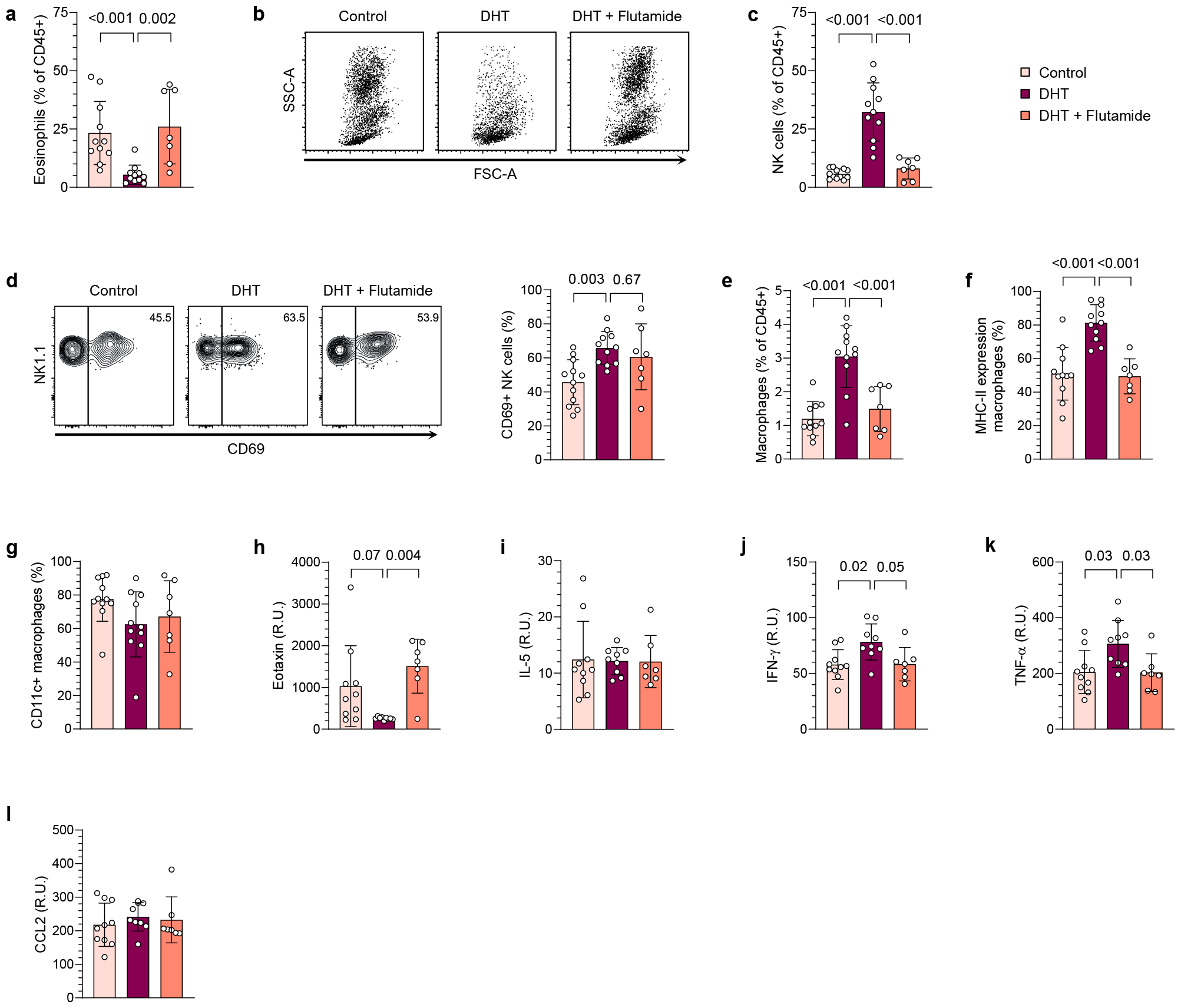
Uterine eosinophil and NK cell populations are markedly altered by androgen exposure in PCOS-like mice. (**a**) Frequency of eosinophils among immune cells in uterus *(n* = 11 Control, 11 DHT, 7 DHT + Flutamide). (**b**) Representative plots of CD45^+^ immune cells, gated on live single cells. (**c**) Frequency of NK cells in uterus *(n* = 12, 11, 7). (**d**) CD69 expression on NK cells in uterus *(n* = 12, 11, 7). (**e**) Frequency of macrophages in uterus *(n* = 11, 11, 7). (**f**) MHC-II expression on macrophages *(n* = 11, 11, 7). (**g**) CD11c expression on macrophages *(n* = 11, 11, 7). (**h**) Eotaxin (CCL11) levels in uterus *(n* = 10, 9, 7). (**i**) IL-5 levels in uterus *(n* = 10, 9, 7). (**j**) IFN-γ levels in uterus *(n* = 10, 9, 7). (**k**) TNF-α levels in uterus *(n* = 10, 9, 7). (**l**) CCL2 levels in uterus *(n* = 10, 9, 7). Data are presented as means ± SD. n indicates the number of biologically independent samples examined. Statistical analysis was assessed by Kruskal-Wallis with Dunn’s multiple comparison (**a, g**), one-way ANOVA with Dunnett’s multiple comparison (**c**-**f**) or mixed-effects ANOVA with Bonferroni’s multiple comparison test (**h**-**l**) and significant differences were indicated with p values. Source data are provided as a Source Data File.

### Ovarian macrophage populations are decreased by androgen exposure in PCOS-like mice

Anovulation and aberrant ovarian steroidogenesis are central components in the pathology of PCOS. Considering the distinct alterations observed in the uterine immune populations, we next investigated whether and how androgen exposure affects immune cells in ovaries, one of the most central organs in PCOS. Flow cytometry analysis revealed a decreased frequency of macrophages in ovaries of DHT-exposed mice (Fig. 3a). NK cells in ovaries of DHT-exposed mice were unchanged in terms of frequencies (Fig. 3b), while a higher proportion expressed CD69, an effect that was prevented by co-treatment with flutamide (Fig. 3c). However, there was an overall increase in CD4^+^ T helper cells in ovaries of DHT-exposed mice compared to controls and mice co-treated with flutamide, an opposite effect to the trend seen in blood, while cytotoxic CD8^+^ T cells were unaltered (Fig. 3d-e). Finally, there was no clear effect on the frequency of neutrophils or eosinophils in ovaries (Supplementary Fig. 3a-b). Taken together, ovarian immune cells are affected by androgen exposure, an effect that is mediated through AR activation.

**Figure 3.**
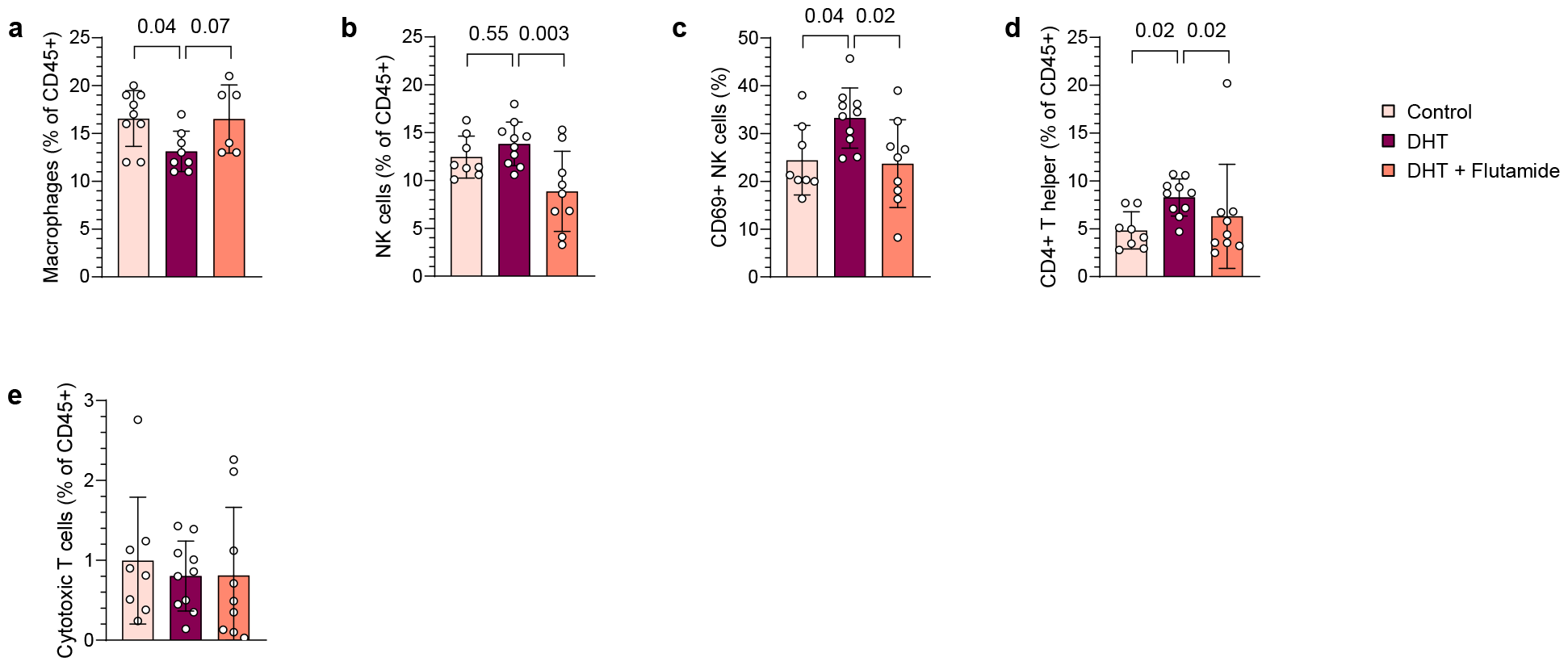
Ovarian macrophage populations are decreased by androgen exposure in PCOS-like mice. (**a**) Frequency of macrophages in ovaries, expressed as percent of CD45^+^ immune cells *(n* = 9 Control, 8 DHT, 6 DHT + Flutamide). (**b**) Frequency of NK cells in ovaries *(n* = 8, 10, 9). (**c**) CD69 expression on NK cells in ovaries *(n* = 8, 10, 9). (**d**) Frequency of CD4^+^ T helper cells in ovaries *(n* = 8, 10, 9). (**e**) Frequency of CD8^+^ cytotoxic T cells in ovaries *(n* = 8, 10, 9). Data are presented as means ± SD. n indicates the number of biologically independent samples examined. Statistical analysis was assessed by one-way ANOVA with Dunnett’s multiple comparison (**a**-**c**), or Kruskal-Wallis with Dunn’s multiple comparison (**d**-**e**) and significant differences were indicated with p values. Source data are provided as a Source Data File.

### The peripubertal DHT-induced mouse model is an insulin-resistant model of PCOS

PCOS is strongly linked to metabolic comorbidities, such as obesity and type-2 diabetes, which have substantial inflammatory components. To assess if these comorbidities are mirrored in our model and to identify any potential confounding factors of obesity on the immune profile, the metabolic phenotype of DHT-exposed mice was investigated. Repeated EchoMRI measurements were performed over 19 weeks following implantation of slow-releasing DHT-pellets to assess if prolonged androgen exposure affects body composition (Fig. 4a). No difference in fat mass was seen between groups at any timepoint (Fig. 4b). However, in line with previous findings^21^, DHT-exposed mice displayed an increased body weight compared to control (Supplementary Fig. 4a), which was due to higher lean mass in DHT-exposed mice (Supplementary Fig. 4b). The prevention of increased lean mass by co-treatment with flutamide was statistically significant at the last timepoint. To rule out that the observations were due to feeding habits, differences in activity, or altered metabolic rate, the mice were analysed in metabolic cages. There were no differences in respiratory exchange ratio, energy expenditure, food intake or locomotor activity between groups (Supplementary Fig. 4c-f). Considering that women with PCOS have a higher risk of type-2 diabetes, independent of BMI^3^, glucose metabolism and insulin response in DHT-exposed mice were assessed by oral glucose tolerance test (oGTT). Glucose uptake did not differ between groups at any time point (Fig. 4c), but DHT-exposed mice displayed an increased insulin response 15 min following glucose administration compared to controls with a concurrent increase in the homeostatic model assessment of insulin resistance (HOMA-IR), indicating insulin resistance (Fig. 4d-e). This effect was prevented by co-treatment with flutamide. Finally, glycated hemoglobin (HbA1c) was analyzed to assess long-term glucose levels in our model. Of note, DHT-exposed mice have a higher HbA1c compared to control and mice co-treatment with flutamide (Fig. 4f), indicating a dysregulated glucose metabolism in these animals. Taken together, the peripubertal DHT-induced mouse model used in this study exhibits an impaired glucose metabolism and insulin response, despite unaltered fat mass.

**Figure 4.**
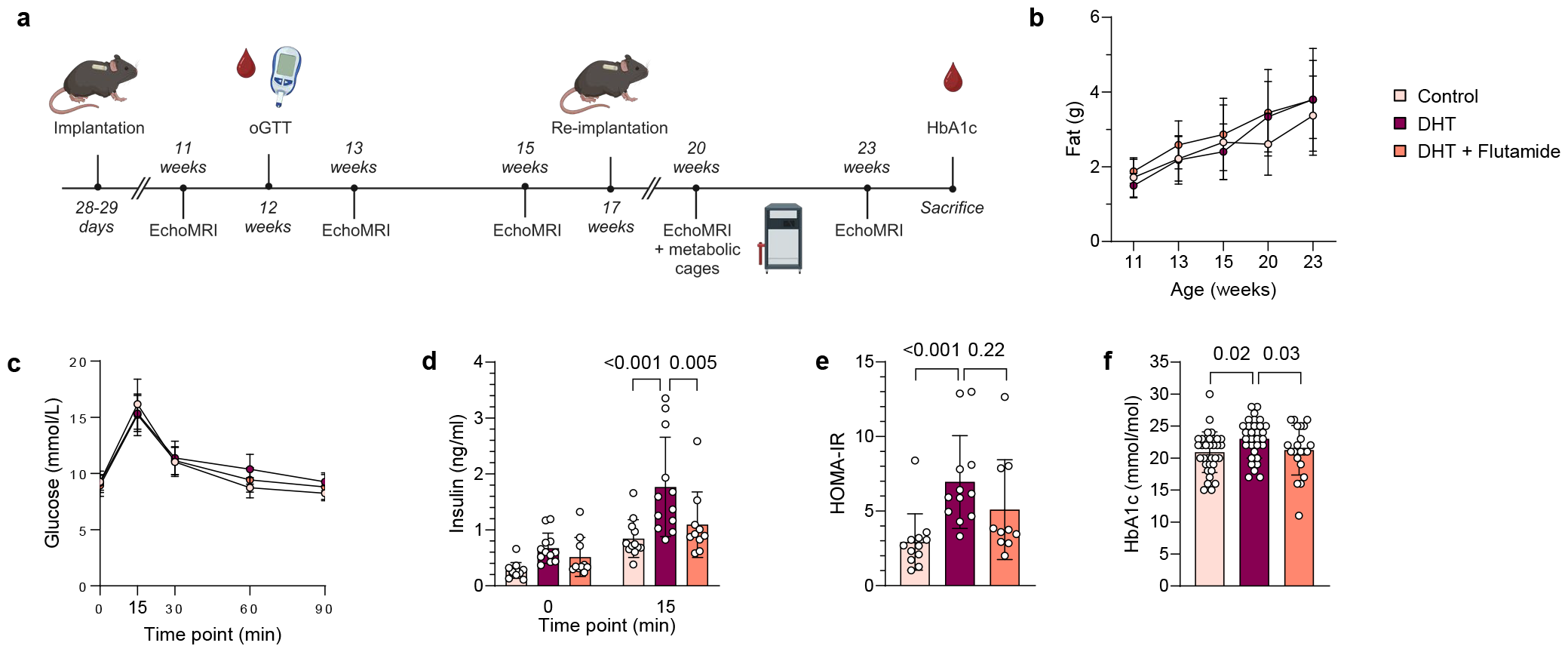
The peripubertal DHT-induced mouse model is a non-obese but insulin resistant model of PCOS. (**a**) Experimental design. (**b**) Fat mass *(n* = 12 Control, 11 DHT, 9 DHT + Flutamide). (**c**) Insulin levels at baseline and 15 min following glucose administration *(n* = 12, 12, 10). (**d**) Blood glucose levels during oGTT *(n* = 12, 12, 10). (**e**) HOMA-IR, calculated from fasted glucose and insulin levels *(n* = 12, 12, 10). (**f**) Glycosylated hemoglobin levels (HbA1c) *(n* = 34, 31, 21). Data are presented as means ± SD. n indicates the number of biologically independent samples examined. Statistical analysis was assessed by two-way ANOVA with Bonferroni’s multiple comparison test (**b**-**d, f**), or Kruskal-Wallis with Dunn’s multiple comparison (**e**) and significant differences were indicated with p values. Source data are provided as a Source Data File.

### DHT-exposed PCOS-like mice display an aberrant immune profile in VAT albeit unaltered fat mass

Type 2 diabetes is prevalent among hyperandrogenic women with PCOS, independent of BMI^3,11^, and immune cells in VAT are known to play a pivotal role in regulating metabolic function^7^. Accordingly, we investigated whether the impaired glucose metabolism and insulin resistance in DHT-exposed mice is mediated by an aberrant immune profile in adipose tissue. To this end, the immune profile in VAT of DHT-induced PCOS-like mice was characterized by flow cytometry. A distinct gating strategy was applied (Supplementary Fig. 5a) as macrophages in VAT displayed distinctly different characteristics compared to those in other analyzed tissues. Likely due to phagocytosis of lipid droplets^22^, macrophages in VAT varied from SSC^low^ to SSC^high^, although predominantly SSC^low/mid^ (Supplementary Fig. 5b). Since macrophages in VAT could not be clearly distinguished from neutrophils based on the expression of Ly6G, macrophages in VAT were defined as CD45^+^CD11b^+^SSC^low/mid^Ly6G^mid/high^F4/80^+^ while neutrophils were defined as CD45^+^CD11b^+^SSC^mid^Ly6G^high^F4/80^-^. Consistent with an unaltered fat mass, the frequency of macrophages was unchanged in VAT of DHT-exposed mice (Fig. 5a). Closer analysis of the macrophage population revealed that these clearly could be divided based on their expression of CD11b and CD11c (Fig. 5b). Interestingly, while no effect was seen on CD11b^mid^ CD11c^-^ macrophages (Supplementary Fig. 5c), the frequency of CD11b^high^ CD11c^+^ macrophages was increased in DHT-exposed mice, which was prevented by co-treatment with flutamide (Fig. 5b). Subsequent analysis showed that the expression of MHC-II differed between these subpopulations, with CD11b^high^ CD11c^+^ macrophages predominantly being MHC-II^mid^ while a larger proportion of CD11b^mid^ CD11c^-^ macrophages display a high expression of MHC-II (Fig. 5c-d). Notably, the expression of MHC-II on CD11b^mid^ CD11c^-^ macrophages differed between groups; DHT-exposed mice displayed a lower frequency of CD11b^mid^ CD11c^-^ MHC-II^high^ macrophages compared to controls. As in the uterus, the eosinophil (CD45^+^CD11b^+^SSC^high^Siglec-F^+^F4/80^-^) population was reduced in VAT of DHT-exposed mice, which was prevented by co-treatment with flutamide (Fig. 5e). In line with the effect observed in blood and ovaries of DHT-exposed mice, a higher frequency of NK cells expressed CD69 in VAT (Fig. 5f), which was reversed in mice co-treated with flutamide, with no effect on the overall frequency of NK cells (Fig. 5g). Consistent with findings in uterus, DHT-exposure did not affect the absolute count of immune cells in VAT (Supplementary Fig. 5d). The frequencies of T cells and neutrophils were also unchanged (Supplementary Fig. 5e-i). Next, cytokine levels in VAT were analyzed by Bio-Plex to assess how androgen exposure affects the cytokine profile. In contrast to uterus, IL-5 was reduced in VAT of DHT-exposed mice, with no clear effect on eotaxin (Fig. 5h-i), suggesting that eosinophils are insufficiently maintained in the tissue. Furthermore, IL-4 and IL-13 were lower in VAT of DHT-exposed mice (Fig. 5j-k), which could be due to the reduced eosinophil population. Opposite to the effect seen in uterus, IFN-γ and TNF-α were found to be lower in VAT of DHT-exposed mice (Fig. 5l-m), possibly contradicting a higher activation state of the NK cells. Interestingly, IL-10 and IL-2, generally regarded as anti-inflammatory cytokines, were also decreased in VAT of DHT-exposed mice (Fig. 5n-o). No difference was seen on IL-1β or GM-CSF (Supplementary Fig. 5j-k). To conclude, these findings demonstrate that androgens induce changes in immune cell composition and cytokine profile in VAT of PCOS-like mice, independent of fat mass.

**Figure 5.**
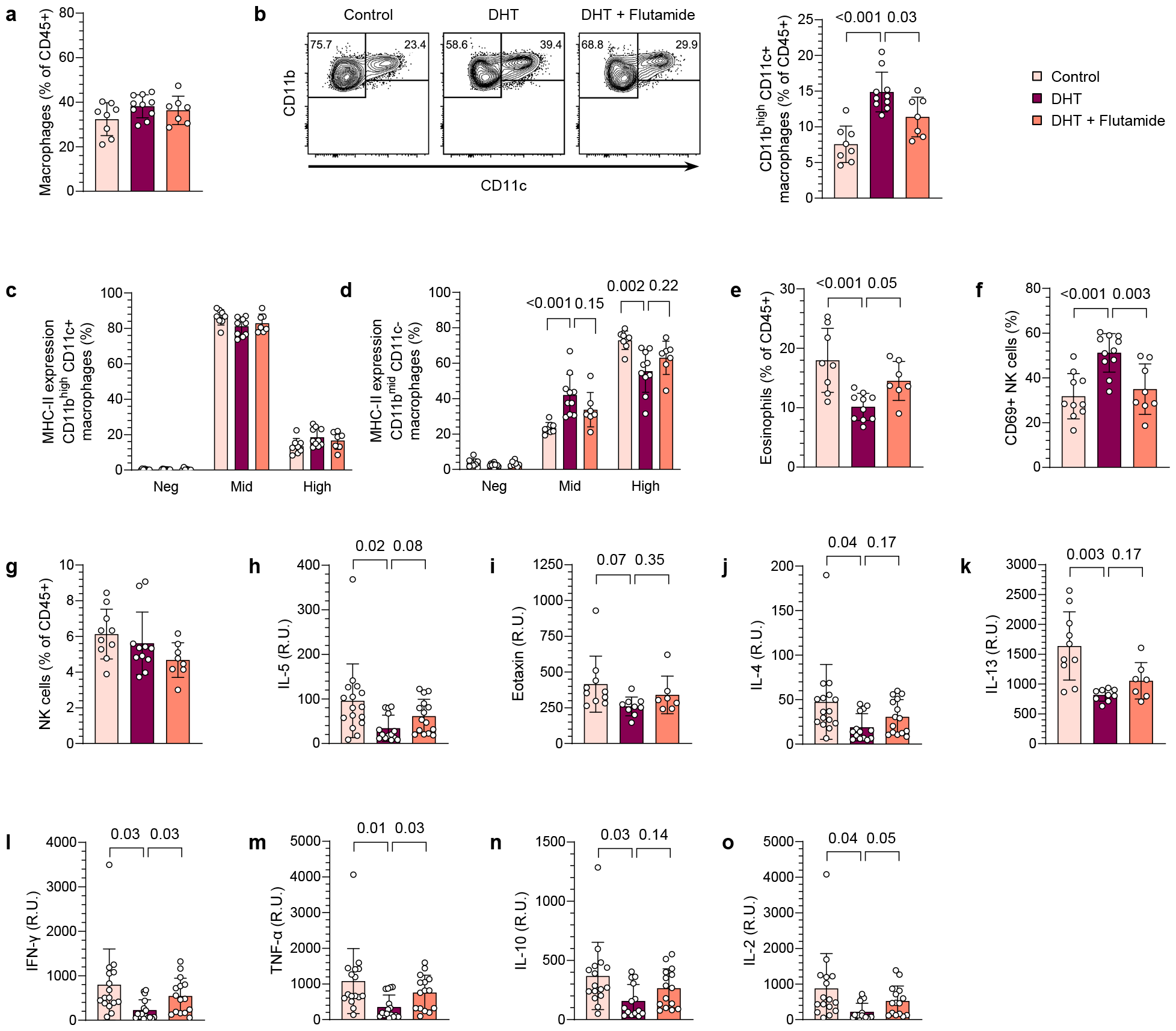
DHT-exposed PCOS-like mice display an aberrant immune profile in VAT albeit unaltered fat mass. (**a**) Frequency of macrophages among immune cells in VAT *(n* = 8 Control, 10 DHT, 7 DHT + Flutamide). (**b**) Representative plots of subpopulations of macrophages (CD45^+^CD11b^+^SSC^low/mid^F4/80^+^Ly6G^mid^) based on expression of CD11b and CD11c and frequency CD11b^high^CD11c^+^. (**c**) MHC-II expression on CD11b^high^CD11c^+^ macrophages *(n* = 8, 10, 7). (**d**) MHC-II expression on CD11b^mid^CD11c^-^ macrophages *(n* = 8, 10, 7). (**e**) Frequency of eosinophils in VAT *(n* = 8, 10, 7). (**f**) CD69 expression on NK cells in VAT *(n* = 10, 11, 8). (**g**) Frequency of NK cells in VAT *(n* = 10, 11, 8). (**h**) IL-5 levels in VAT *(n* = 16, 14, 15). (**i**) Eotaxin (CCL11) levels in VAT *(n* = 10, 9, 7). (**j**) IL-4 levels in VAT *(n* = 16, 14, 15). (**k**) IL-13 levels in VAT *(n* = 10, 9, 7). (**l**) IFN-γ levels in VAT *(n* = 16, 16, 15). (**m**) TNF-α levels in VAT *(n* = 16, 16, 15). (**n**) IL-10 levels in VAT *(n* = 16, 14, 15). (**o**) IL-2 levels in VAT *(n* = 16, 16, 15). Data are presented as means ± SD. n indicates the number of biologically independent samples examined. Statistical analysis was assessed by one-way ANOVA with Dunnett’s multiple comparison (**a**-**f**), Kruskal-Wallis with Dunn’s multiple comparison (**g**), or mixed-effects ANOVA with Bonferroni’s multiple comparison test (**h**-**o**) and significant differences were indicated with p values. Source data are provided as a Source Data File.

## Discussion

Mounting evidence points to a low-grade inflammation in PCOS pathology, although conflicting reports and improperly powered studies render the effect of hyperandrogenism on circulating and tissue-specific immune populations inconclusive^13,15,16,18,19,23–26^. Utilizing a well-established peripubertal PCOS-like mouse model induced by continuous DHT exposure^20^, we demonstrate that androgen exposure impacts circulating and tissue resident immune cells differently. These findings underscore the significance of distinguishing between circulating and tissue-resident immune cells and align with existing studies demonstrating phenotypic and functional differences among immune populations within these compartments^27–31^. Importantly, the co-treatment with flutamide, an AR antagonist used for the treatment of hyperandrogenism and hirsutism in women with PCOS^11^, enabled us to conclude that these effects on the immune cells were dependent on AR activation. Since mature eosinophils and NK cells do not express the AR^32,33^, the modulations of these populations were likely mediated through indirect mechanisms, such as AR signaling on stroma cells or other leukocytes, or a maintained effect from AR expressing progenitors.

We consistently observed a higher proportion of CD69^+^ NK cells in all examined tissues of mice exposed to androgens. CD69 is considered both a marker of activation and tissue-residency^28^, suggesting a higher activation state of circulating NK cells in DHT-exposed mice. However, it is important to note that tissue-resident NK cells exhibit distinct phenotypes compared to their circulating counterparts^27^. The higher frequency of NK cells expressing CD69 in our PCOS-like mice may thus indicate a specific impact on the tissue-resident populations. Indeed, this observation is substantiated by the augmented presence of NK cells in the uterus of DHT-exposed mice. While the higher levels of IFN-γ and TNF-α in uteri could suggest an excessively activated state, it is important to note that both are essential factors for endometrial function; IFN-γ secreted by uNK cells is crucial for remodeling of spiral arteries and TNF-α is an essential mediator during embryo implantation^34^. These observed alterations of NK cells in our model are particularly intriguing given the suggested endometrial dysfunction in women with PCOS. Moreover, we see a drastic decrease in the number of eosinophils, which normally constitute the dominant population in the uterus of mice during diestrus^35^. As eotaxin is required for homing of eosinophils into the uterus^36^, the lower levels seen in our model clearly indicate that the reduced eosinophil population is due to a disrupted recruitment from the periphery. DHT-exposed mice also display an increased frequency of macrophages, which is in line with a recent report claiming that the frequencies of macrophages are higher in endometrium of women with PCOS compared to BMI-matched controls^23^. The increased number of macrophages in our model does not appear to be due to a higher infiltration of monocytes that differentiate into macrophages as the frequency of monocytes in the uterus was unchanged between groups and is further supported by unaltered levels of chemoattractant CCL2. How these modulations affect endometrial vascularization and embryo implantation calls for further investigations.

Anovulation and aberrant ovarian steroidogenesis represent pivotal components in the pathology of PCOS. It is well-established that immune cells play an important role in ovarian function^5,37,38^. Macrophages promote ovulation through the secretion of cytokines and matrix metalloproteinases^5^. The decreased frequency of macrophages in ovaries of PCOS-like mice is therefore intriguing. However, whether this contributes to ovarian dysfunction, or is a mere consequence of anovulation due to androgen exposure, requires further elucidation.

In contrast to uterus, the overall frequency of NK cells remained unchanged in VAT. Nevertheless, the higher expression of CD69 on NK cells in VAT of DHT-exposed mice is interesting as tissue-resident NK cells in adipose tissue of mice fed high-fat diet can induce insulin resistance by promoting the differentiation of macrophages towards a pro-inflammatory phenotype through IFN-γ^39^. Although the overall levels of IFN-γ and TNF-α were lower in VAT of DHT-exposed mice, we did not specifically determine the ability of NK cells to release IFN-γ. Therefore, the possibility of a higher activation state of tissue-resident NK cells cannot definitively be ruled out. This aspect merits further investigation, particularly considering that the phenotype of macrophages was indeed altered in the VAT of DHT-exposed mice. Macrophages residing in adipose tissue play a central role in the regulation of glucose homeostasis and are known to accumulate in the adipose tissue of individuals with obesity^7^.

Consistent with the unaltered fat mass in our study, the overall frequency of macrophages remained unchanged in VAT of DHT-exposed mice. However, specific subpopulations of macrophages were distinctly affected by androgen exposure. These subpopulations were clearly discernible based on their expression of CD11c and CD11b. Notably, the frequency of CD11b^high^ CD11c^+^ macrophages were exclusively increased in DHT-exposed mice. Furthermore, androgens affected the expression of MHC-II on CD11b^mid^ CD11c^-^ but not CD11b^high^ CD11c^+^ macrophages. These results are interesting as an enhanced pro-inflammatory phenotype of adipose tissue macrophages is detrimental for metabolic function and contributes to insulin resistance^7^. Indeed, DHT-exposed mice displayed a mild insulin resistance, as evidenced by elevated HbA1c, increased HOMA-IR, and a higher insulin response following glucose challenge during oGTT. Contrary, macrophages with an anti-inflammatory phenotype are essential for maintaining insulin sensitivity in mice with diet-induced obesity^9^. Eosinophils have been shown to sustain the anti-inflammatory phenotype of macrophages through the release of IL-4 and IL-13^10^. Here we observed a lower frequency of eosinophils as well as reduced levels of IL-4 and IL-13 in VAT of DHT-exposed mice, which could potentially influence the function of macrophages in adipose tissue. In contrast to uterus, levels of eotaxin in VAT were unchanged, while IL-5 was lower. This suggests that the decreased eosinophil population in VAT may be attributed to a lack of survival factors or local proliferation rather than a disruption in recruitment from the periphery. The intriguing association between an aberrant immune profile in adipose tissue and insulin resistance is particularly fascinating considering the unaltered fat mass observed in this study. This observation holds significance in light that women with PCOS have adipose tissue dysfunction^26^ and a higher prevalence of type 2 diabetes independent of BMI^3,11^. Whether this aberrant immune profile drives the metabolic dysfunction observed in DHT-exposed mice remains to be elucidated.

Strikingly in blood, DHT-exposure induced an overall reduction in immune cells, without a clear reduction in any specific population. Nevertheless, a trend suggested a reduced population of CD3^+^ T cells. This coupled with a decreased thymic weight could suggest a potential effect of androgens on thymopoiesis, as previously described in male mice^40^. In contrast to circulating immune cells, no discernible effect of androgens was observed on the overall numbers of CD45^+^ immune cells in any of the investigated tissues.

In conclusion, we demonstrate that hyperandrogenism causes distinct alterations of immune cells in reproductive and metabolic tissues, which could have considerable implications on tissue function and contribute to the reduced fertility and metabolic comorbidities associated with PCOS.

## Methods

### Animals and model induction

23 days old female C57BL/6JRj were purchased from Janvier Labs. Mice were kept under a 12-hour light/dark cycle in a temperature-controlled environment with access to water and normal chow diet (CRM (P), Special Diets Services) ad libitum. For model induction, continuously releasing DHT implants were made as previously described^41^. Briefly, silastic tubes were filled with ∼2.5 mg DHT (5α-Androstan-17β-ol-3-one, ≥97.5%, Sigma-Aldrich) and trimmed to a total length of 5 mm. 28-29 days old mice received implants subcutaneously in the neck region under anaesthesia with 1.5-2% isoflurane (Vetflurane, Virbac). Control mice were implanted with an empty, blank implant. To investigate AR activation, a third group was implanted with both DHT implant and a 25 mg continuously releasing flutamide pellet (NA-152, Innovative Research of America). Implants and flutamide pellets were replaced 90 days after implantation by the same procedure. All animal experiments were approved by the Stockholm Ethical Committee for animal research (20485-2020) in accordance with the Swedish Board of Agriculture’s regulations and recommendations (SJVFS 2019:9) and the European Parliament’s directive on the protection of animals used for scientific purposes (2010/63/EU). Animal care and procedures were performed in accordance with the guidelines by the Federation of European Laboratory Animal Science Associations (FELASA) and controlled by Comparative Medicine Biomedicum at the Karolinska Institutet in Stockholm, Sweden.

### Tissue collection

Estrous cyclicity was assessed by vaginal smears and determined by vaginal cytology as previously described^42^. Mice were sacrificed at diestrus at age 13-14 or 24-26 weeks. Blood was collected by cardiac puncture under anesthesia with 3% isoflurane (Vetflurane, Virbac) into EDTA-coated tubes (Sarstedt) and kept on ice. Plasma was separated by centrifugation at 2000 rcf for 10 min at 4°C and stored at –80°C. Organs were harvested and wet weight registered. For flow cytometric analysis, perigonadal VAT, uterus and ovaries were collected in RPMI-1640 (Sigma-Aldrich) supplemented with 2% fetal bovine serum (FBS, Gibco™), retroperitoneal lymph nodes and 3 mesenteric lymph nodes and half the spleen, cut sagittal, were collected in Ca^2+^ and Mg^2+^ free DPBS (Sigma-Aldrich). For cytokine analysis, VAT and uterus were snap frozen in liquid nitrogen and stored at –80°C.

### Flow cytometry

Uteri and ovaries were minced using fine scissors and digested by collagenase type I (1 mg/ml, Sigma-Aldrich) and 0.8U DNase I (Sigma-Aldrich) in RPMI-1640 (Sigma-Aldrich) with 2% FBS (Gibco™), whilst gently shaken at 37°C for 20 and 15 min respectively. VAT was minced using fine scissors and digested by collagenase type IV (1 mg/ml, Sigma-Aldrich) in RPMI-1640 (Sigma-Aldrich) with 2% FBS (Gibco™), whilst gently shaken at 37°C for 20-30 min. Samples were passed through a 100μm strainer to obtain a single cell suspension. Lymph nodes and spleen were passed directly through a 100μm strainer. Cells were washed in flow cytometry buffer (2% FBS and 2mM EDTA in PBS). Erythrocytes in blood, VAT and spleen were hemolyzed (0.16M NH_4_Cl, 0.13M EDTA and 12mM NaHCO_3_ in H_2_O) followed by additional wash. Following incubation with Fc-block (anti-mouse CD16/32, clone 2.4G2, BD Biosciences), cell surface markers were detected using fluorochrome-conjugated antibodies: CD45 (30-F11, Invitrogen/ eBioscience), CD11b (M1/70, BD Biosciences), Siglec-F (E50-2440, BD Biosciences), Ly6G (1A8, BD Biosciences), Ly6C (AL-21, BD Biosciences), F4/80 (BM8, eBioscience), MHC II (M5/114.15.2, eBioscience), CD11c (N418, BD Biosciences), CD3e (500A2, BD Biosciences), NK-1.1 (PK136, BD Biosciences), CD8a (53-6.7, BD Biosciences), CD4 (RM4-5, BD Biosciences), CD25 (PC61, BD Biosciences), CD69 (H1.2F3, eBioscience). LIVE/DEAD Fixable Aqua/Far Red Dead Cell stain (Molecular Probes, Invitrogen) was added for exclusion of nonviable cells. Cells were analyzed with BD^®^ LSR II SORP Flow Cytometer (BD Biosciences). Data was further analyzed by FlowJo™ v10.8 Software (BD Life Sciences).

### Bio-Plex

Cytokines in VAT (GM-CSF, IFN-γ, IL-1β, IL-2, IL-4, IL-5, IL-10, TNF-α) were analyzed using Bio-Plex Pro Mouse Cytokine 8-plex Assay (Bio-Rad) according to the manufacturer’s instructions. Cytokines in uterus and VAT (IL-1α, IL-1β, IL-2, IL-3, IL-4, IL-5, IL-6, IL-9, IL-10, IL-12 (p40), IL-12 (p70), IL-13, IL-17A, Eotaxin, G-CSF, GM-CSF, IFN-γ, KC, MCP-1 (MCAF), MIP-1α, MIP-1β, RANTES, TNF-α) were analyzed using Bio-Plex Pro Mouse Cytokine 23-plex Assay (Bio-Rad). Briefly, frozen tissue was processed using Bio-Plex® Cell Lysis Kit (Bio-Rad) according to manufacturer’s instructions. Protein concentrations were determined by Pierce BCA (ThermoFisher) and diluted in cell lysis buffer to 200–900 μg/ml. Samples were stored at –20°C until analysis. Cytokine levels were normalized to protein concentrations for tissue samples. Analytes that were below detection level were excluded from the analysis.

### Metabolic phenotyping

Body weight was recorded weekly. Body composition was measured biweekly in conscious mice by EchoMRI (Echo-MRI-100™, EchoMRI LLC, Houston, TX, US) from age 11 weeks throughout the study (Fig. 4a). Glucose metabolism was assessed by oGTT at age 12 weeks. Mice were fasted for 5 hours in clean cages. D-glucose was administered by oral gavage (2 mg/g of body weight) and blood glucose was measured at baseline and 15, 30, 60 and 90 min using a glucose meter (Free Style Precision). For insulin measurements, blood was collected into EDTA-coated tubes by tail bleeding at baseline and 15 min. Plasma was separated by centrifugation at 2000 rcf for 10 min at 4°C and stored at –20°C. Insulin was measured by ELISA (Crystal Chem). Metabolic activity was assessed by indirect calorimetry (TSE systems) with concurrent recordings of food intake and spontaneous locomotor activity. 20 weeks old mice were kept individually (N = 5-6 per group) in metabolic cages for three consecutive days. The first 24 hours were considered acclimatization period and not analyzed. Glycosylated hemoglobin levels (HbA1c) were measured by DCA® HbA1c Reagent Kit (Siemens Healthcare) on a DCA Vantage® Analyzer (Siemens Healthcare) at the time of sacrifice.

### Statistical analysis

No statistical methods were used to predetermine sample size, based on previous reported assessments^43^. Animals were arbitrarily allocated to experimental groups without formal randomization. Investigators were not formally blinded to group allocation during the experiment. Indirect calorimetry was analyzed using CalR^44^, data are presented as mean ± SEM. Differences between groups were determined by ANCOVA with body mass as covariate. All other data are presented as mean ± SD, one animal considered an experimental unit. Normality was assessed by Shapiro-Wilk test. For normally distributed data, differences between groups were determined by one-way ANOVA with Dunnett’s multiple comparison test, or by two-way ANOVA and Bonferroni’s multiple comparison test when measurements were repeated. Data that didn’t follow a normal distribution were analyzed by Kruskal-Wallis with Dunn’s multiple comparison test. For HbA1c, three cohorts were pooled and analyzed with 2-way-ANOVA, corrected for age and batch variability. Cytokine levels were analyzed by mixed-effects ANOVA with Geisser-Greenhouse correction for unequal variability and Bonferroni’s multiple comparison test. P ≤ 0.05 was considered statistically significant. Statistical analyses were performed using GraphPad Prism 8 (GraphPad Software).

## Data availability

The data that support the findings of this study are available within the paper in source data are provided in BioStudies and will be publicly available upon publication of this paper under accession number: TMP_1706540120334.

## Author contribution

Conceptualization: ST; AA; ESV. Methodology: ST; AA; ESV; MHJ. Investigation: ST; AA; SR; HL; AZ; EL; GE; MB. Data acquisition, analysis, and visualization: ST; AE; MHJ; ESV. Project administration: ST; ESV. Supervision: ESV; AB; MCIK; CS. Wrote the manuscript: ST; ESV. All authors: reviewing and editing manuscript. Funding acquisition: ESV.

## Acknowledgement

We thank the Metabolic Phenotyping Centre at the Strategic Research program in Diabetes at the Karolinska Institutet for the use of the TSE Systems and EchoMRI, and the Biomedicum Flow cytometry Core facility (Karolinska Institutet), supported by KI/SLL, for providing technical expertise and scientific input. This work is supported by Swedish Medical Research Council: project no. 2022-00550 (ESV); Distinguished Investigator Grant – Endocrinology and Metabolism, Novo Nordisk Foundation: NNF22OC0072904 (ESV); Diabetes Foundation: DIA2021-633 (ESV); Karolinska Institutet KID funding: 2020-00990 (ESV).

