## Supplementary information for "Androgens modulate the immune profile in a mouse model of polycystic ovary syndrome"

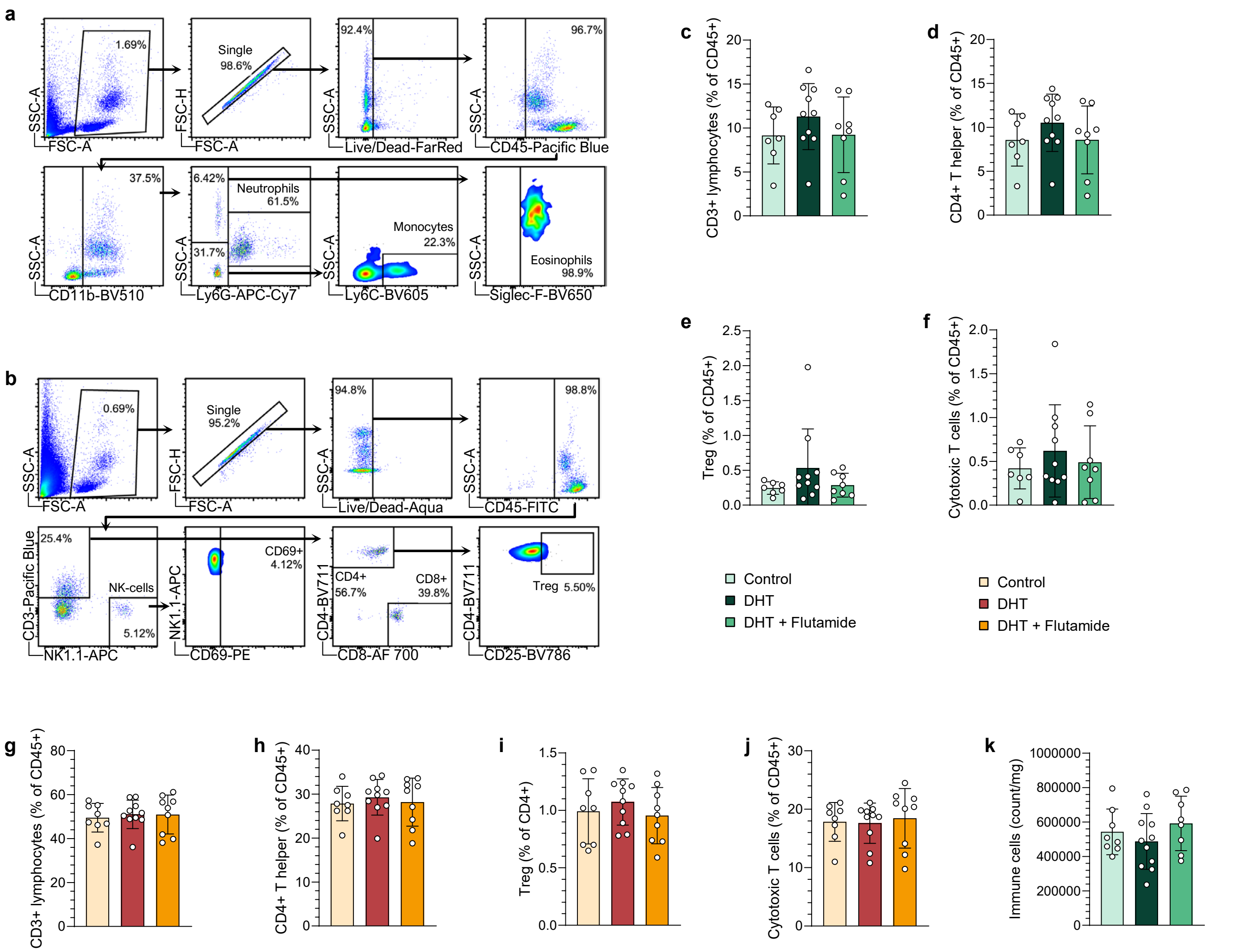


**Supplementary figure 1. Immune populations in blood and secondary lymphoid organs are differently affected by androgen exposure.** (**a**) Gating strategy myeloid cells in blood. (**b**) Gating strategy NK- and T cells in blood. (**c**) Frequency of CD3⁺ T cells in spleen (*n* = 7 Control, 10 DHT, 8 DHT + Flutamide). (**d**) Frequency of CD4⁺ T helper cells in spleen (*n* = 7, 10, 8). (**e**) Frequency of CD4⁺ CD25⁺ T reg cells in spleen (*n* = 7, 10, 8). (**f**) Frequency of CD8⁺ cytotoxic T cells in spleen (*n* = 7, 10, 8). (**g**) Frequency of CD3⁺ T cells in lymph nodes (*n* = 8, 10, 9). (**h**) Frequency of CD4⁺ T helper cells in lymph nodes (*n* = 8, 10, 9). (**i**) Frequency of CD4⁺ CD25⁺ T reg cells in lymph nodes (*n* = 8, 10, 9). (**j**) Frequency of CD8⁺ cytotoxic T cells in lymph nodes (*n* = 8, 10, 9). (**k**) Immune cell count in spleen (*n* = 8, 11, 8). Data are presented as means ± SD. n indicates the number of biologically independent samples examined. Statistical analysis was assessed by one-way ANOVA with Dunnett's multiple comparison (**c**-**d**, **f**-**i**, **k**) or by Kruskal-Wallis with Dunn's multiple comparison (**e**, **j**) Source data are provided as a Source Data File.

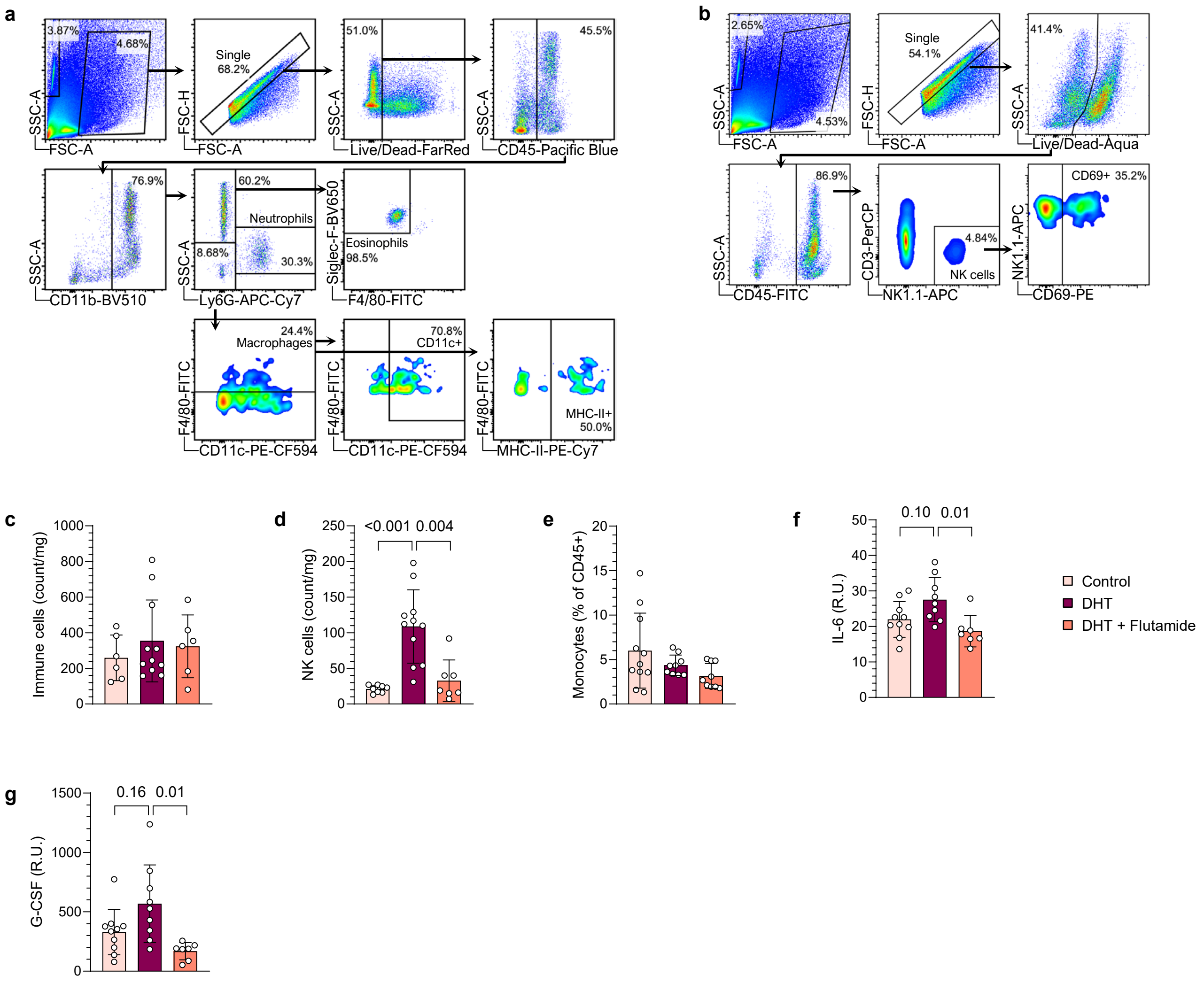


**Supplementary figure 2. Uterine eosinophil and NK cell populations are markedly altered by androgen exposure in PCOS-like mice.** (**a**) Gating strategy myeloid cells. (**b**) Gating strategy NK cells in uterus. (**c**) Number of CD45+ immune cells in uterus *(n* = 6 Control, 11 DHT, 6 DHT + Flutamide). (**d**) NK cell count uterus *(n* = 8, 11, 7). (**e**) Frequency of monocytes in uterus *(n* = 11, 9, 9). (**f**) IL-6 levels in uterus *(n* = 10, 9, 7). (**g**) Granulocyte-colony stimulating factor (G-CSF) levels in uterus *(n* = 10, 9, 7). Data are presented as means ± SD. n indicates the number of biologically independent samples examined. Statistical analysis was assessed by Kruskal-Wallis with Dunn's multiple comparison (**c-e**), or mixed-effects ANOVA with Bonferroni's multiple comparison test (**f-g**) and significant differences were indicated with p values. Source data are provided as a Source Data File.

| Analyte | Comparison | 95,00% CI | Summary | Adjusted P Value |
|---|---|---|---|---|
| IL-1a | DHT vs. Control | -2,54 to 29,3 | ns | 0,11 |
| | DHT vs. Flutamide | -15,1 to 36,4 | ns | 0,58 |
| IL-1b | DHT vs. Control | -0,961 to 3,48 | ns | 0,36 |
| | DHT vs. Flutamide | -2,80 to 4,00 | ns | >0,99 |
| IL-2 | DHT vs. Control | -2,03 to 45,3 | ns | 0,08 |
| | DHT vs. Flutamide | -12,0 to 48,3 | ns | 0,29 |
| IL-3 | DHT vs. Control | -1,77 to 2,60 | ns | >0,99 |
| | DHT vs. Flutamide | -1,53 to 2,97 | ns | 0,86 |
| IL-4 | DHT vs. Control | -1,76 to 4,23 | ns | 0,64 |
| | DHT vs. Flutamide | -1,50 to 4,23 | ns | 0,49 |
| IL-9 | DHT vs. Control | -2,71 to 33,6 | ns | 0,1 |
| | DHT vs. Flutamide | -5,92 to 38,1 | ns | 0,17 |
| IL-10 | DHT vs. Control | -13,6 to 20,8 | ns | >0,99 |
| | DHT vs. Flutamide | -20,0 to 23,7 | ns | >0,99 |
| IL-12p40 | DHT vs. Control | -6,13 to 65,4 | ns | 0,11 |
| | DHT vs. Flutamide | -34,2 to 70,4 | ns | 0,79 |
| IL-12p70 | DHT vs. Control | -34,9 to 133 | ns | 0,34 |
| | DHT vs. Flutamide | -27,3 to 136 | ns | 0,23 |
| IL-13 | DHT vs. Control | -109 to 199 | ns | 0,95 |
| | DHT vs. Flutamide | -77,8 to 240 | ns | 0,42 |
| IL-17a | DHT vs. Control | -2,57 to 3,91 | ns | >0,99 |
| | DHT vs. Flutamide | -4,44 to 5,54 | ns | >0,99 |
| GM-CSF | DHT vs. Control | -12,3 to 29,4 | ns | 0,65 |
| | DHT vs. Flutamide | -13,0 to 27,1 | ns | 0,78 |
| KC | DHT vs. Control | -7,85 to 32,5 | ns | 0,3 |
| | DHT vs. Flutamide | -23,2 to 35,1 | ns | >0,99 |
| MIP-1a | DHT vs. Control | -9,91 to 2,35 | ns | 0,3 |
| | DHT vs. Flutamide | -11,6 to 1,33 | ns | 0,13 |
| MIP-1b | DHT vs. Control | -97,3 to 12,4 | ns | 0,15 |
| | DHT vs. Flutamide | -67,2 to 14,3 | ns | 0,25 |
| RANTES | DHT vs. Control | -198 to 370 | ns | 0,91 |
| | DHT vs. Flutamide | -108 to 458 | ns | 0,26 |

**Supplementary table 1.** Cytokine levels in uterus *(n* = 10 Control, 9 DHT, 7 DHT + Flutamide). Data are presented as 95% confidence intervals (CI) with adjusted p-values. Statistical analysis was assessed by mixed-effects ANOVA with Bonferroni's multiple comparison test. Source data are provided as a Source Data File. ns indicates non-significant.

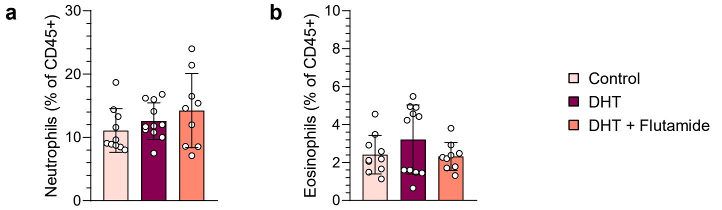


**Supplementary figure 3. Ovarian macrophage populations are decreased by androgen exposure in PCOS-like mice.** (**a**) Frequency of neutrophils in ovaries, expressed as percent of $CD45^{+}$ immune cells (*n* = 10 Control, 11 DHT, 9 DHT + Flutamide). (**b**) Frequency of eosinophils in ovaries (*n* = 10, 11, 9). Data are presented as means ± SD. n indicates the number of biologically independent samples examined. Statistical analysis was assessed by Kruskal-Wallis with Dunn's multiple comparison. Source data are provided as a Source Data File.

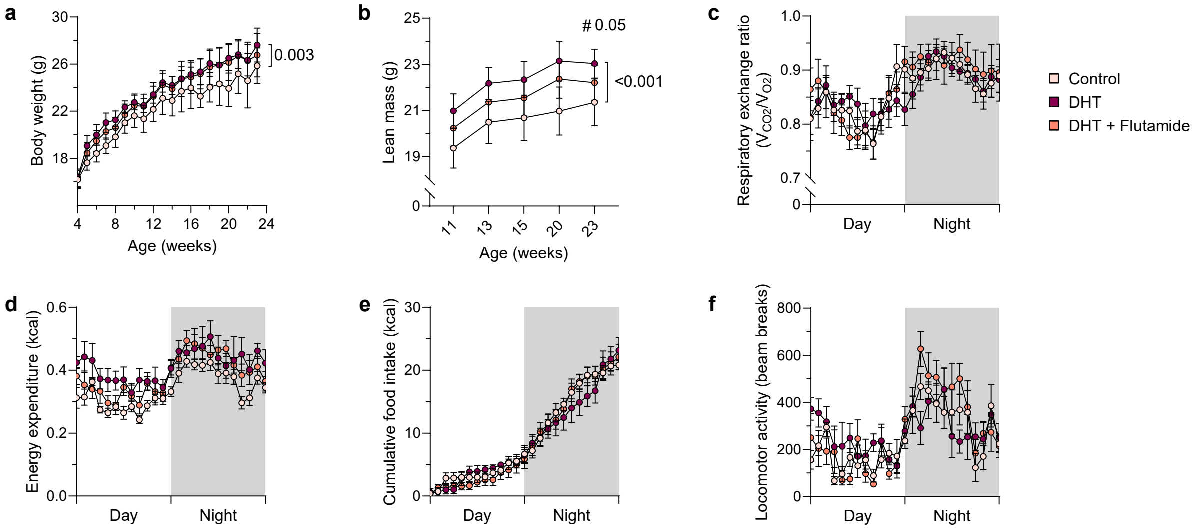


**Supplementary figure 4. The peripubertal DHT-induced mouse model is a non-obese but insulin resistant model of PCOS.** (**a**) Body weight (*n* = 12 Control, 11 DHT, 8 DHT + Flutamide). (**b**) Lean mass (*n* = 12, 11, 8). (**c**) Respiratory exchange ratio (*n* = 6, 5, 5). (**d**) Energy expenditure (*n* = 6, 5, 5). (**e**) Food intake (*n* = 6, 5, 5). (**f**) Locomotor activity (*n* = 6, 5, 5). Data are presented as means ± SD (**a-b**) and means ± SEM (**c-f**). # indicates statistical significance between DHT and DHT + Flutamide. n indicates the number of biologically independent samples examined. Statistical analysis was assessed by mixed-effects ANOVA with Bonferroni's multiple comparison test (**a**), two-way ANOVA with Bonferroni's multiple comparison test (**b**), or ANCOVA with body mass as covariate (**c-f**) and significant differences were indicated with p values. Source data are provided as a Source Data File.

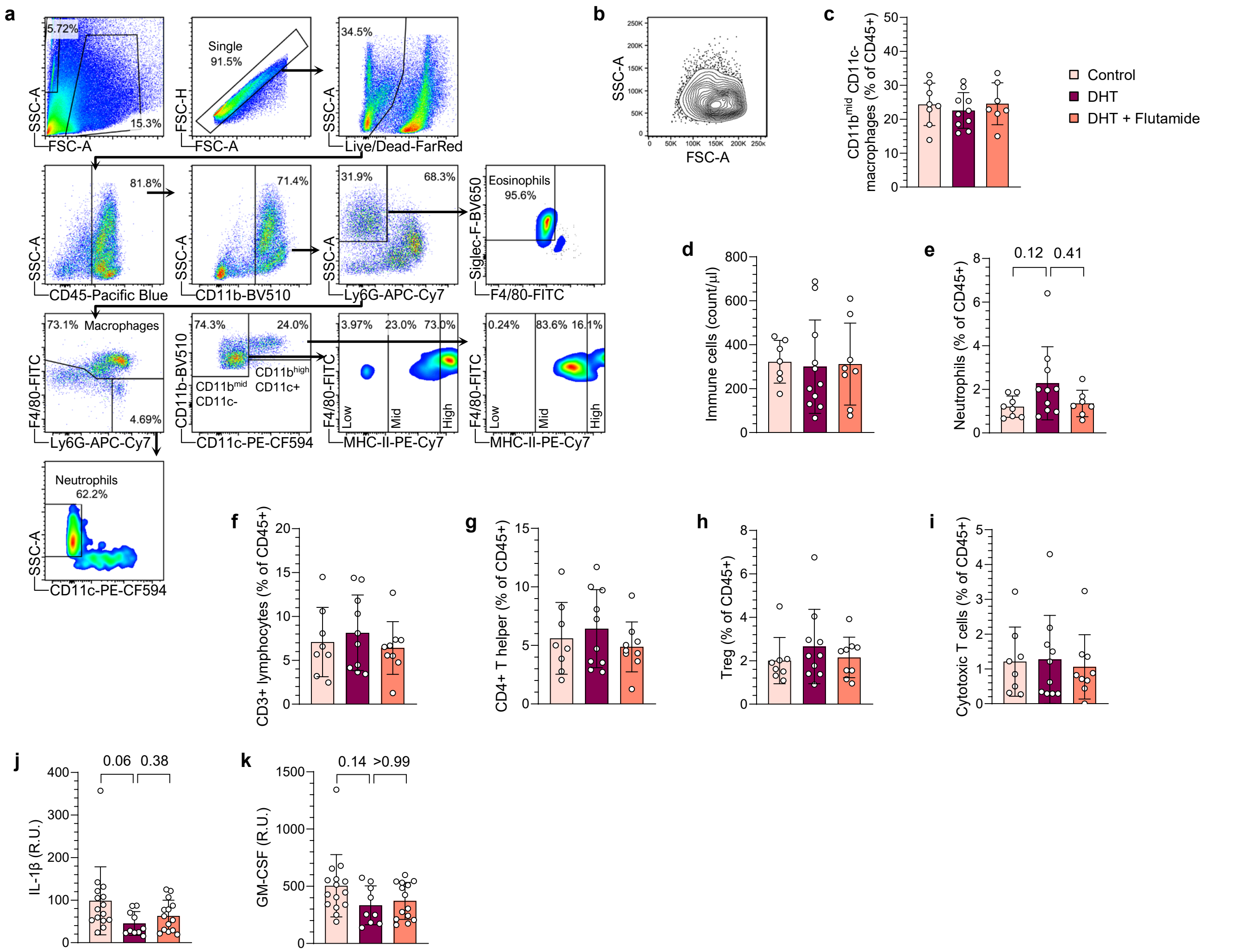


**Supplementary figure 5. DHT-exposed PCOS-like mice display an aberrant immune profile in VAT albeit unaltered fat mass.** (**a**) Gating strategy for myeloid cells in VAT. (**b**) Representative plot of macrophages in VAT. (**c**) Frequency CD11b$^{mid}$CD11c$^{-}$ macrophages in VAT (*n* = 8 Control, 10 DHT, 7 DHT + Flutamide). (**d**) Number of CD45+ immune cells in VAT (*n* = 7, 11, 8). (**e**) Frequency of neutrophils in VAT (*n* = 8, 10, 7). (**f**) Frequency of CD3$^{+}$ T cells in VAT (*n* = 8, 10, 9). (**g**) Frequency of CD4$^{+}$ T helper cells in VAT (*n* = 8, 10, 9). (**h**) Frequency of CD4$^{+}$ CD25$^{+}$ T reg cells in VAT (*n* = 8, 10, 9). (**i**) Frequency of CD8$^{+}$ cytotoxic T cells in VAT (*n* = 8, 10, 9). (**j**) IL-1β levels in VAT (*n* = 15, 10, 14). (**k**) GM-CSF levels in VAT (*n* = 15, 9, 14). Data are presented as means ± SD. n indicates the number of biologically independent samples examined. Statistical analysis was assessed by one-way ANOVA with Dunnett's multiple comparison (**c-d**, **f-g**), Kruskal-Wallis with Dunn's multiple comparison (**e**, **h-i**), or mixed-effects ANOVA with Bonferroni's multiple comparison test (**j-k**) and significant differences were indicated with p values. Source data are provided as a Source Data File.
